# Comprehensive Mutational Landscape Analysis of Monkeypox Virus Proteome

**DOI:** 10.1101/2024.09.19.613877

**Authors:** Tugba Ozaktas, Ayten Dizkirici Tekpinar, Alessandra Carbone, Mustafa Tekpinar

## Abstract

In this study, we present a comprehensive computational analysis of the single point mutational landscapes of the Monkeypox virus (MPXV) proteome. We reconstructed full single-point mutational landscapes of 171 MPXV proteins using an advanced mutational effect predictor, ESCOTT, selected for its superior performance on viral proteins. ESCOTT performance was assessed by benchmarking against the experimental data in the ProteinGym (v1.0.0) dataset that contains 48917 multiple and 173502 single point mutations. A recent MPXV strain sequenced in July 2024 was used as the reference genome. Multiple sequence alignments and protein structures were generated using Colabfold v1.5.5, and the predicted structures were evaluated with pLDDT metric, secondary structure predictions, and comparisons with available experimental data, ensuring high confidence in the structural models. We determined mutational sensitivity of all positions in a protein utilizing ESCOTT scores and demonstrated their functional implications on cysteine proteinase and helicase of MPXV. Moreover, we created an interactive visualization tool to visualize mutational landscapes and sensitivities in a publicly available Google Colab. Furthermore, we introduced a novel, interpretable metric (Average Gene Mutation Sensitivity) to prioritize the most mutation-sensitive proteins within the large MPXV proteome as prime candidates for drug or vaccine development. Among the top 20 proteins identified with this metric, several were membrane-associated proteins, proven to be important for viral interactions with the hosts in other viruses. This analysis provides a valuable resource for assessing the impact of new MPXV variants. This pioneering study underscores the significance of understanding MPXV evolution in the context of the ongoing global health crisis and offers a robust computational framework to support this effort.

## Introduction

World Health Organization declared Mpox outbreak a public health emergency of international concern on August 14, 2024. Data between January 1, 2024 and January 16, 2025 shows 24872 confirmed cases with 90 deaths (http://www.cdc.gov/mpox/situation-summary/index.html?cove-tab=0). Understanding impact of mutations in MPXV is of utmost importance both for assessing evolution of the virus and development of novel vaccines or therapies against this emerging health threat. Unfortunately, there are no systematic investigations of variant effects with experimental methods like deep mutational scanning. Therefore, the computational prediction of the single point mutational landscape of the MPXV genome represents a critical frontier in understanding the virus’s evolution, epidemiology, and potential responses to therapeutic interventions.

The MPXV has a large double-stranded DNA genome of approximately 197 kb in size, encoding around 180 proteins (1, 2). Compared to other orthopoxviruses, the MPXV genome is larger, with a higher number of open reading frames (3). Despite the fact that the mutational landscape of MPXV is characterized by a relatively slow mutation rate compared to RNA viruses, significant variations have been observed in recent strains, particularly those emerging post-2022. This discrepancy underscores the importance of ongoing surveillance and genomic analysis to monitor the virus’s evolution and potential adaptive changes that could affect public health responses (4, 5). Researchers can better understand the dynamics of MPXV and inform strategies for vaccine development and therapeutic interventions by utilizing computational mutational impact prediction tools. The computational prediction of MPXV’s mutational landscape not only improves our understanding of the virus’s evolutionary patterns but also serves as a vital resource for public health initiatives aimed at controlling future outbreaks.

Due to these reasons, we first determined the best computational variant prediction algorithm for viruses utilizing the viral proteins in a publicly available experimental dataset called ProteinGym (v1.0.0). Our study revealed that ESCOTT (6) method is the best mutational effect prediction algorithms for viral proteins. We generated input multiple sequence alignments (MSA) and structure files with Colabfold v1.5.5. We assessed structure qualities with respect to a few available experimental structures and evaluated structure qualities over the entire genome with the pLDDT (predicted Local Distance Difference Test) (7, 8) and secondary structure analysis. Then, we calculated complete single point mutational landscapes of 171 MPXV proteins with ESCOTT using Colabfold generated MSAs and structure files. We predicted mutational sensitivity of each position in a protein with a quantity called Average Minmaxed Mutation Score, derived from ESCOTT scores. Finally, we analyzed sensitivity of each gene to mutations with a metric called ‘Average Gene Mutation Sensitivity’ and identified the top 20 MPXV genes. We demonstrated that AGMS analysis can help to prioritize target proteins for therapeutic development.

## Methods

### ESCOTT Calculations for Viral Proteins in ProteinGym

We obtained multiple sequence alignment (MSA) files and protein structures using Colabfold v1.5.5 (9, 10) because Colabfold produces smaller MSA files with sufficient diversity in a time-efficient manner. Since ESCOTT requires protein structures as well as MSA files, AlphaFold generated PDB files provided by ProteinGym v1.0.0 were used as structural inputs of ESCOTT.

### Reference MPXV Genome

Since we wanted to use a recent strain of the MPXV, we selected "http://www.ncbi.nlm.nih.gov/nuccore/PQ008840.1, which has been sequenced in July, 2024. We generated fasta files for 175 proteins. OPG003, OPG025, OPG108, OPG147 genes were eliminated from the sequence set due to long unknown segments. The list of the 171 remaining proteins is given in Table S1 of Supplementary Information.

### MSA and Structure Files for MPXV Proteins

In consistence with ProteinGym MSA file generation procedure, we obtained MSA files and protein structures using Colabfold method via its web server (9, 10). We removed the gaps in query sequences of the MSA files and converted them from a3m format to fasta format with reformat utility tool of HHSuite (11). Number of sequences in each MSA file is given in Table S1 of the Supplementary Information. We visualized 3D structures of the proteins with Pymol (12). Secondary structure calculations were performed with biotite Python library and dssp (13, 14). Colorblind friendly viridis colormap for Pymol (15) was obtained from https://github.com/smsaladi/pymol_viridis and AlphaFold (10) pLDDT colormap for Pymol comes from https://github.com/BobSchiffrin/pymol_scripts/blob/main/colour_af_plddt.py.

### Calculation of Full Single Point Mutational Landscapes with ESCOTT and iGEMME

We used PRESCOTT program to calculate ESCOTT and iGEMME scores of all single point amino acid mutations of 171 genes (6, 16). iGEMME is an improved version of GEMME utilizing only evolutionary information to determine impact of mutations while ESCOTT uses both evolutionary and structural information for the same purpose (6, 16). The docker image containing version 1.6.0 of PRESCOTT at https://hub.docker.com/r/tekpinar/prescott-docker was used both for ESCOTT and iGEMME computations (6). All details about ESCOTT method can be found at (6).

ESCOTT raw scores can be between [-12, 2], where low scores indicate high mutational impact, and high scores no effect. The raw scores are not uniform across different proteins. Due to this reason, we sort all scores in the ESCOTT/iGEMME matrix by their rank and divide the rank position by the total number of mutations in the matrix to obtain a score between zero and one. Since we want low scores (close to zero) to indicate no mutational impact and high scores (close to one) suggest high mutational impact, we subtract the final scores from 1. All rank-sorted ESCOTT/iGEMME scores are obtained using this approach.

### Average Minmaxed Mutation Score (AMMS)

Sometimes, it is necessary to measure sensitivity of a position to mutations regardless of the identity of a target mutation. Here, we suggest a metric to quantify this positional sensitivity with a parameter called Average Minmaxed Mutation Score (AMMS). To achieve our purpose, we applied the following procedure: Raw ESCOTT or iGEMME scores are minmax normalized to obtain values between zero and one. All values are subtracted from one in order to make zero values to indicate no effect and values close to one to indicate high impact upon mutation. Then, Average Minmaxed Mutation Score (AMMS) is calculated by summing up all mutation scores divided by 20 for each amino acid position in a protein.

### Average Gene Mutation Sensitivity (AGMS)

As explained above, AMMS is obtained to measure positional sensitivity to mutations within a protein. Here, we used AMMS to calculate AGMS, a per protein quantity, for a protein with N amino acids as follow:

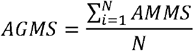

AGMS is a score between zero and one that can help prioritize drug or vaccine targets. Low values suggest low sensitivity to mutations for the gene, while scores close to one indicate high sensitivity to mutations for the gene. It should be noted that if ‘rank-sorted’ values had been used in AMMS, AGMS would always be 0.5 and that’s why we used minmax normalized AMMS scores to calculate AGMS.

## Results and Discussions

### Selecting the Best Mutational Effect Prediction Algorithm for Viral Proteins

Extended ProteinGym dataset (v1.0.0) contains 217 deep mutational scanning substitution experiments from various organisms (17). Moreover, the dataset contains theoretical predictions from 62 methods for all of the experimental measurements. ProteinGym v1.0.0 includes 28 virus experiments containing 48917 multiple and 173502 single point mutations (Table S2). Of these 28 experiments, only three of them (highlighted with red text color in Table S2) contain multiple mutations. When Spearman correlation performance is ordered by taxon, GEMME (16) and TranseptEVE family of models (L, M and S models) (18) are the best performers for mutational effect prediction of viral proteins (see the benchmarks at https://proteingym.org/benchmarks for the performance of the other 58 methods). In addition to the forementioned methods, we investigated performances of two recently developed methods called ESCOTT and iGEMME. We calculated all mutations for 28 viral proteins with these methods using the Colabfold MSA files and structure files provided by ProteinGym. Our results show that ESCOTT is the best method both for predicting effect of combined mutations (single+multiple mutations) (Fig. 1a) and single point mutations only (Fig. 1b), while GEMME (16) becomes the second-best method. When only multiple point mutations from 3 proteins are considered, ESCOTT and iGEMME are the top predictors (Fig. 1c). In addition to mutation type analysis, we wanted to see if prediction quality changes depending on genetic material of the virus. There are 24 proteins from RNA viruses and 4 proteins from DNA viruses (see Table S2 for details). ESCOTT is the best prediction method both for RNA and DNA virus proteins (Fig. 1d and e). While GEMME becomes the second method for RNA viruses, iGEMME is the second-best method for DNA virus proteins. In terms of computational performance, ESCOTT is very fast and it can be run on CPUs compared to TranceptEVE requiring a recent generation of GPU.

**Fig. 1.**
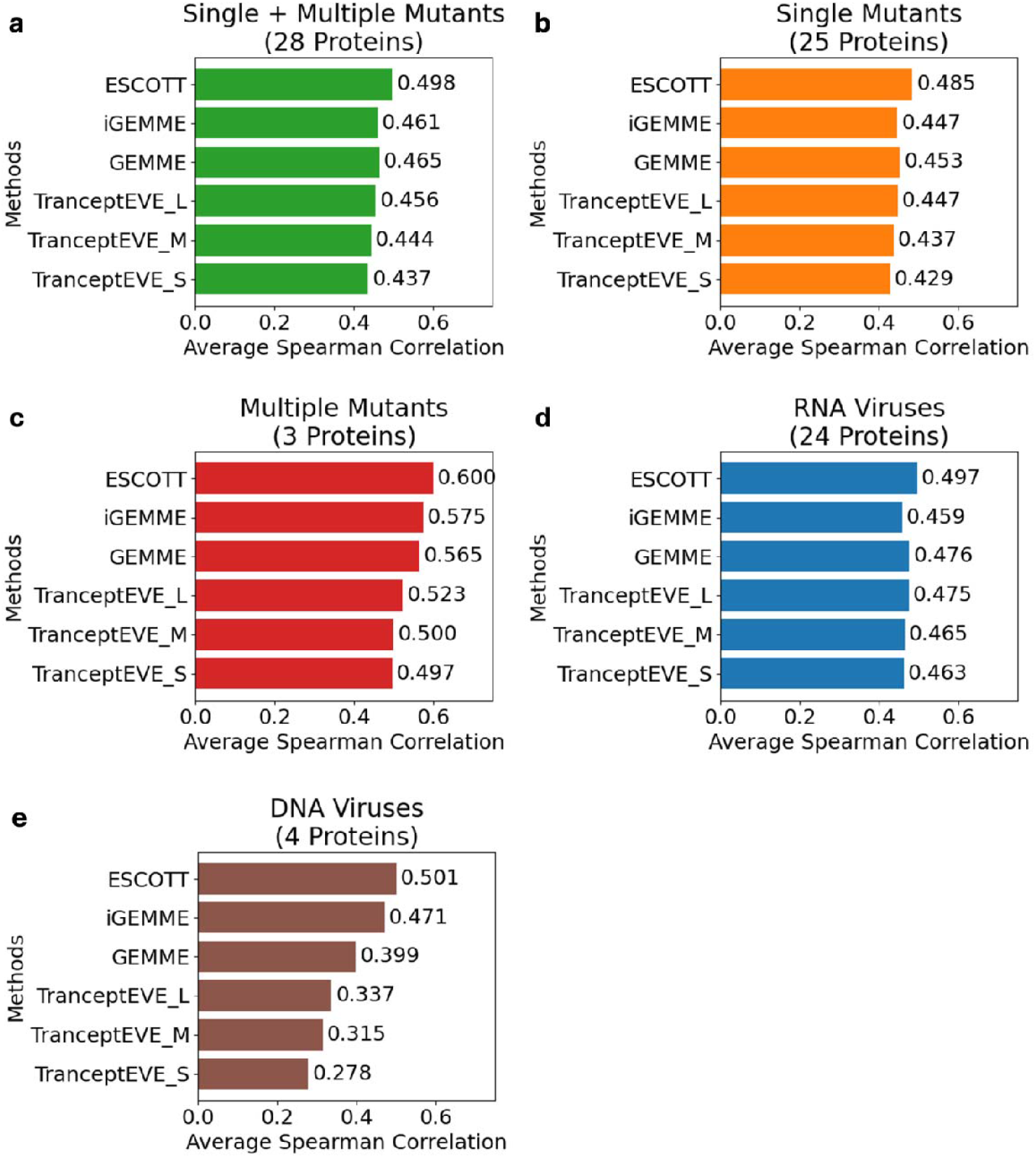
Performance comparison of ESCOTT with the other five methods. MSA files generated by Colabfold were used for ESCOTT and iGEMME calculations. Number of proteins in each dataset is given in parenthesis under the title of the subfigures. a) Average Spearman correlation performance of the best six methods over 28 deep mutational scanning experiments of viral proteins from ProteinGym v1.0.0 dataset containing both single and multiple point mutations. b) Average Spearman correlation performance of the best six methods over 25 deep mutational scanning experiments of viral proteins from the same dataset containing only single point mutations. c) Average Spearman correlation performance of the best six methods over 3 deep mutational scanning experiments of viral proteins containing only multiple point mutations. d) Average Spearman correlation performance for proteins obtained from 24 RNA viruses. e) Average Spearman correlation performance for proteins obtained from 4 DNA viruses.

We observed no relationship between the number of sequences in the MSA and prediction performance of ESCOTT (see Fig. S1A and B). For instance, the ENV_HV1BR protein MSA contained 11,044 sequences yet had a modest Spearman correlation of 0.352, whereas the POLG_PESV protein, with only 210 sequences, achieved a higher correlation of 0.554. These findings clearly show that MSA depth does not determine ESCOTT prediction performance.

Finally, we wanted to see ESCOTT and iGEMME performances using MSA files provided by ProteinGym. Our results show that ESCOTT is the best method when multiple and single point mutations are evaluated together (Fig. S2A). ESCOTT is again the best algorithm predicting mutational effects both for single point mutations only and multiple point mutations only (Fig. S2B and C). If viruses are classified according their genetic material, ESCOTT remains as the top-performing method (Fig. S2D and E). On the other hand, iGEMME becomes the second method consistently when ProteinGym-provided MSA files are used both for RNA and DNA viral proteins.

### Quality of MPXV Protein Structures for Better ESCOTT Predictions

ESCOTT utilizes structural information coming from PDB format files in addition to evolutionary information coming from an MSA file. Since quality of structure prediction may affect the mutational landscape calculations, it is important to check it for better ESCOTT predictions. Due to this reason, we investigated if there are experimental structures of MPXV proteins. We obtained experimental structures of A42R profilin-like protein (PDB ID: 4qwo) (19) and DNA polymerase (PDB ID: 8wpe) (20) of MPXV from Protein Databank (21). We superimposed them with the Colabfold predicted structures of OPG171 and OPG071 using Pymol, respectively (Fig. 2a and b). RMSD was 0.36 Å for A42R profilin-like protein and it was 1.89 Å for DNA polymerase, which suggests a good prediction. pLDDT is another metric used to check the quality of predictions (7, 8). When we inspect pLDDT values projected onto these two structures, we can see that majority of pLDDT values are greater than 90 (shown as dark blue), indicating a good prediction. Since we wanted to find the quality of structure prediction for 171 proteins, we collected pLDDT for all amino acids in all of the predicted structures and plotted them as a pie chart (Fig. 2c). We observed that 75.5% of pLDDT values are very high or high (dark blue or light blue). Moreover, we investigated structure quality from secondary structure perspective. As a result, we investigated if the predicted protein structures contain coils in a large percentage, which is an indicator of disorder in a protein. To do this, we determined secondary structure types of all amino acids and plotted their distributions as a pie chart (Fig. 2d). We observed that coils constitute only 24.9% of the secondary structure types. In conclusion, RMSD comparison with available experimental structures, pLDDT values and secondary structure analysis of the predicted structures suggest that the quality of the predicted structures is good for utilization in mutational effect predictions of all proteins encoded by MPXV genome.

**Fig. 2.**
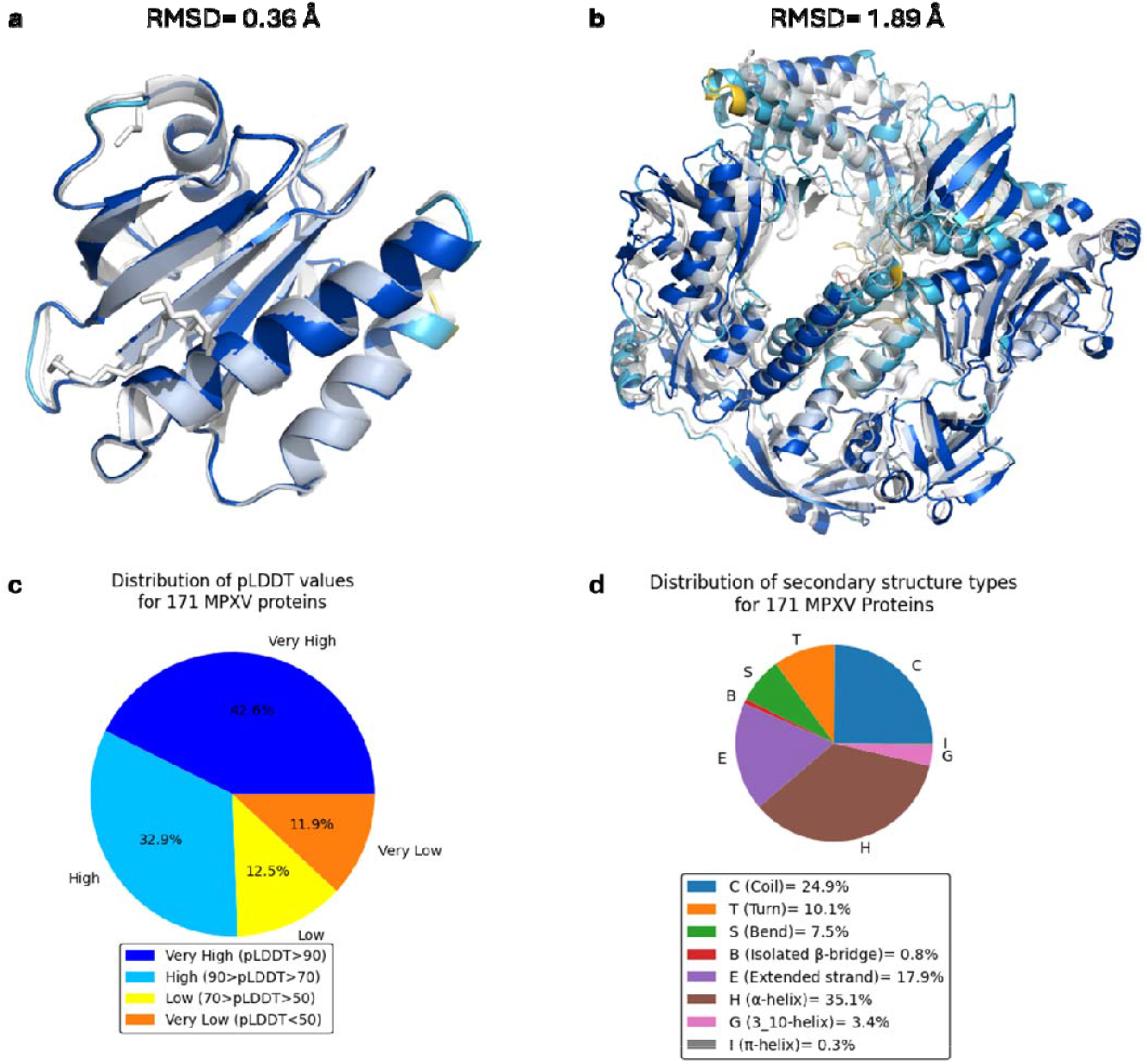
Protein structure quality investigation with RMSD, pLDDT and secondary structure analysis. a) Experimental structure (PDB ID: 4qwo, transparent white cartoon representation) vs Colabfold generated OPG171 structure for profilin domain protein b) Experimental structure (PDB ID: 8wpe, transparent white cartoon representation) vs Colabfold generated OPG071 structure for MPXV DNA polymerase c) Pie chart for distribution of pLDDT values of 171 MPXV proteins. d) Pie chart for distribution of secondary structures for 171 MPXV proteins.

### Single Point Mutational Landscapes of MPXV Proteins and A Graphical User Interface for Their Visualizations

We calculated single point mutational landscapes of 171 MPXV proteins with ESCOTT method. Raw and rank-sorted score files as well as mutational landscapes in PNG image format for all proteins can be downloaded using publicly available zenodo repository from https://doi.org/10.5281/zenodo.13365666.

Since it can be cumbersome for experimental community to extract and visualize data from text files, we provide an interactive, intuitive and publicly available Google Colab for visualization and analysis of all mutational landscapes with CC-BY-NC-SA 4.0 license, which is a license that promotes code and data sharing for the benefit of all scientific community. The Google Colab file publicly is accessible at https://colab.research.google.com/github/tozaktas/mpxv-mutations/blob/master/monkeypox_mutations.ipynb. Moreover, the end-users can download the raw and ranksorted ESCOTT/iGEMME output files for a selected protein from the zenodo repository given above and upload it to PRESCOTT web server (http://prescott.lcqb.upmc.fr/visualisation.php) for an interactive visualization of the results.

### Functional Implications of ESCOTT Predictions over Two Examples

#### i. MPXV Cysteine Proteinase Analysis

As an example of the ESCOTT output, we present here the rank-sorted impact of mutations to all possible amino acids for virus core cysteine proteinase, whose main role is to degrade peptides or proteins (Fig. 3a). Also, we provide the raw scores obtained with ESCOTT in Fig. S3. In addition to full mutational landscapes, we calculated AMMS for each position and added them to occupancy column of a PDB file for all proteins as explained in Average Minmaxed Mutation Score (AMMS) subsection of the Methods section. We projected these per residue sensitivities onto the proteinase structure (see Fig. 3a for AMMS strips and Fig. 3b for the projections). In many viral proteinases, there is a catalytic dyad (or triad) that function like a chemical scissor (22). Catalytic dyad of this proteinase is H241 and C328. It is clear both from the dark colors in Fig. 3b and the ESCOTT AMMS scores given in Fig. 3a and c that the catalytic dyad amino acids are among the most sensitive positions to mutations. Furthermore, we highlighted Calpha atoms of four residues (317, 329, 369 and 373) as spheres, which has AMMS values greater than 0.875. Residues 317 and 329 are in neighborhood of the catalytic dyad. Even tough residues 369 and 373 are not in close proximity of the catalytic dyad, they are also highly buried residues. These results imply that ESCOTT scores and AMMS scores (derived from the ESCOTT scores) can identify correctly functionally important locations in proteins.

**Fig. 3.**
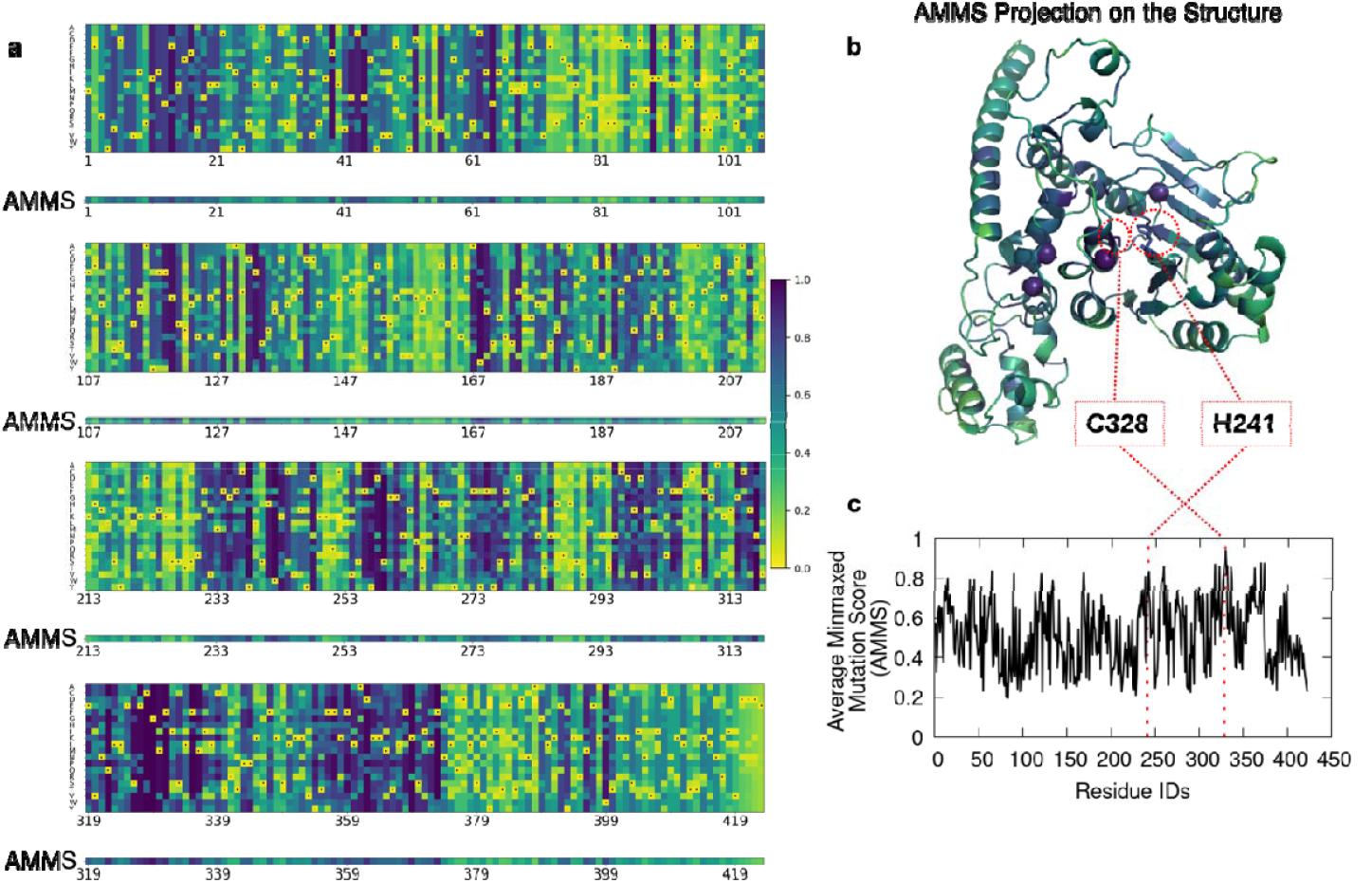
Mutational landscape of virus core cysteine proteinase (OPG083) and its projections obtained with ESCOTT. a) Single point mutational landscape of virus core cysteine proteinase (OPG083) calculated with ESCOTT is given in rank-sorted scale (0, 1) and viridis colormap. Average Minmaxed Mutation Score (AMMS) is given as a strip below each position. Yellow color (values close to zero) indicates no effect while dark blue (values close to one) indicates a high effect for the mutations. Dotted squares denote the original amino acid in the protein sequence. b) AMMS values projected onto Colabfold generated protein structure (OPG083). Catalytic dyad (H241 and C328) of the proteinase is shown in licorice representation in red dotted circles and it has a dark color in viridis colormap, suggesting high mutational sensitivity of the positions. Calpha atoms of four residues (317, 329, 369 and 373) with AMMS values greater than 0.875 are highlighted as spheres. c) 2D plot of the AMMS vs residue IDs. Locations of H214 and C328 are highlighted with red vertical dashed lines.

#### ii. MPXV Helicase Analysis

Helicase is a protein that unwinds double stranded DNA/RNA and it is important in replication of many viruses including MPXV (23, 24). A recent experimental study investigated impact of 9 mutations on DNA unwinding capacity of MPXV helicase and reported three measurements for each mutation as fraction of unwound DNA (25). Even though ESCOTT was not developed to predict amount of functional activity in any protein, we extracted ESCOTT rank-sorted scores for those 9 mutations and compared them with the experimental predictions. Keeping in mind that 0.0 ESCOTT_rank-sorted_ score indicates ‘no impact’ and 1.0 score indicates impactful mutations, we subtracted the scores from 1.0 to make them compatible with experimental measurements. Five mutations (F630A, K509A, N605A, S510A, R620A) impair DNA unwinding almost completely and ESCOTT predicts 4 of them almost perfectly (Fig. 4a). ESCOTT predicts that S510A mutation may impair the function partially while the experiment reports a full disfunction for this mutation. When we investigate location of S510 on experimental structure, we observe that S510 coordinates with Mg^2+^, which will promote the hydrolysis of the cofactor ATP (Fig. 4b) (25). ESCOTT predicts a low activity for L655A mutation while experimental data suggests an activity around 40%. Side chain of this residue forms extensive stacking interactions with the nucleobase of ATP (25). Source of these partially correct predictions is potentially due to the fact that ESCOTT model does not account for interactions of ions, ligands, nucleic acids and other proteins with the protein in question.

**Fig. 4.**
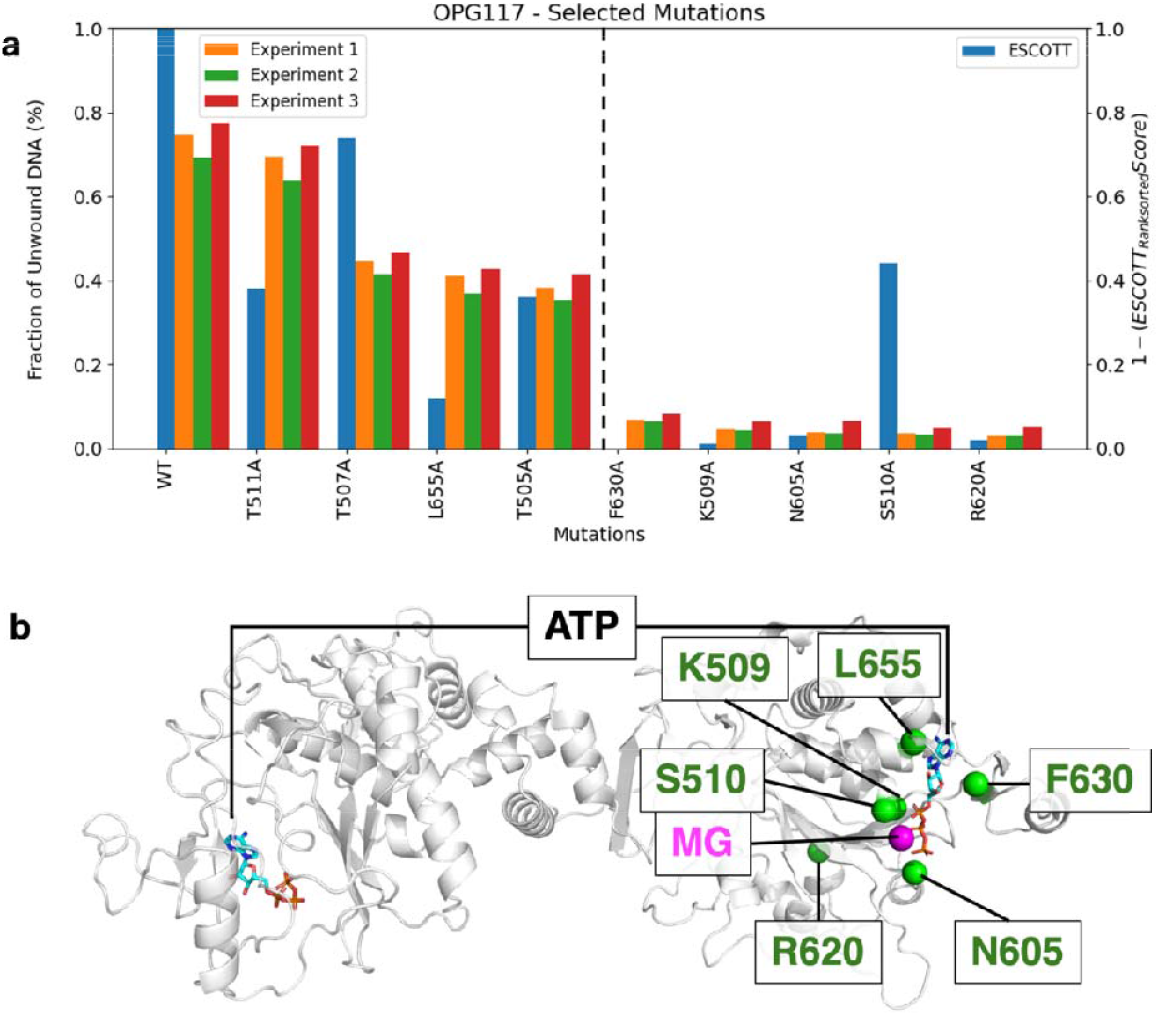
Experimentally measured MPXV helicase activity vs. ESCOTT scores for the wild type and 9 mutations. a) Fraction of unwound DNA measured by three experiments (orange, green and red bars) and rank-sorted ESCOTT scores (blue bars) for the wild type and 9 mutations. Five mutations causing significant disruptions to the helicase activity are on the right-hand side of the plot. b) Locations of ATPs, Mg ion and five disruptive mutations on chain A of experimental structure (PDB ID: 8hwa) are highlighted on gray cartoon representation of the protein. ATP is in licorice representation while Mg ion is shown as a magenta sphere. Calpha atoms of the five disruptive mutations are shown as green spheres.

### MPXV Proteome Analysis with a New Parameter for Target Prioritization

We developed a ranking method to identify viral proteins that may be more sensitive to mutational changes and thus serve as promising targets. To achieve this, we calculate a straightforward measure -the Average Gene Mutation Score (AGMS)-using ESCOTT results from 171 proteins, and then select the top 20 genes with the highest scores (Fig. 5a). The highest-ranking protein, OPG076, is a key component of the entry-fusion complex (EFC). Its conservation across all chordopoxviruses and essential role in viral entry have been well documented (26), and studies have shown that viruses lacking EFC proteins exhibit markedly reduced infectivity and impaired membrane fusion (27). The second highest-ranking gene, OPG059, encodes Cytochrome C oxidase. A recent influenza virus study underscores its role in viral replication (28). Next, OPG158 (A32.5L) encodes a protein similar to the vaccinia virus A32 protein, which is involved in virion membrane assembly (29, 30). Research by Cassetti et al. demonstrated that repression of the A32 gene results in the production of noninfectious, DNA-deficient, spherical, enveloped particles in vaccinia virus (31). Moreover, another study observed that a vaccinia virus strain lacking the B14R ortholog (analogous to A32.5L) displayed reduced virulence, suggesting that A32.5L may similarly impact monkeypox virus pathogenesis, depending on the infection route (29). In addition, the F14 protein from MPXV (OPG058) exhibits 98.6% sequence identity to its vaccinia virus ortholog (UniProt ID: P68707, PG058_VACCW), and Albarnaz et al. demonstrated that F14 promotes virulence in vaccinia virus—a role that is likely conserved in variola, monkeypox, and cowpox viruses (32).

**Fig. 5.**
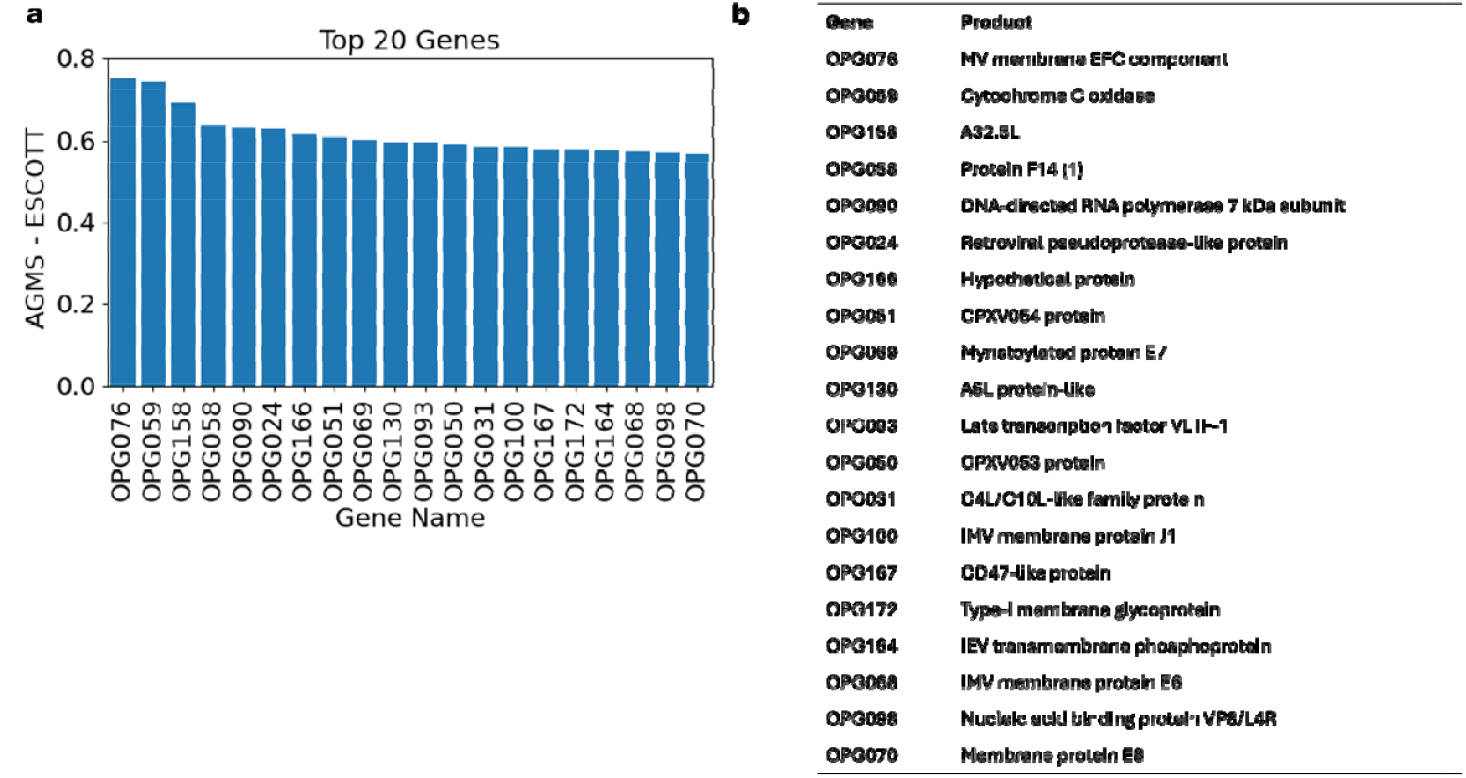
a) Top 20 genes with the highest average gene mutation sensitivity (AGMS) obtained from ESCOTT calculations. b) Gene names and their products for the top 20 genes.

Recognizing that viral membranes are crucial interfaces for host interactions, we also focused on membrane proteins, which are prime drug targets (33, 34). Our AGMS analysis identified seven membrane proteins (OPG076, OPG100, OPG172, OPG164, OPG166, OPG068, and OPG070) (Fig. 5b). For example, OPG100 in vaccinia virus (J1R) is essential for viral growth (35, 36), while a homolog of OPG164 (A36R) is involved in virion transport to the host cell surface (37). Additionally, the 567-amino acid OPG068 protein of MPXV, with 98.9% sequence identity to its vaccinia counterpart, plays a vital role in immature virion formation (38, 39). Furthermore, we examined whether protein length influences AGMS. Our analyses revealed only a weak correlation, with Spearman correlation coefficients of –0.134 for ESCOTT data (Fig. S4). This indicates that AGMS is effectively independent of protein length. Finally, we provide AGMS values for all 171 proteins in Fig. S5 for further inquiry. In summary, our AGMS-based ranking method, robustly identifies a set of viral proteins with fundamental functions and potential as novel drug targets.

## Conclusions

We calculated entire single point mutational landscape of 171 MPXV genes using the state-of-the art ESCOTT method. We provide a Google Colab to visualize the mutational landscapes and AMMS scores on protein structures. ESCOTT mutational landscapes can be used to asses potential impact of new MPXV protein variants while AMMS scores can be used to assess functionally important regions within a protein. Furthermore, we calculated AGMS for all genes in the dataset. AGMS values can also be quite useful for drug and vaccine design studies since the proteins with high AGMS values can be used to prioritize potential drug/vaccine targets.

ESCOTT utilizes evolutionary, physico-chemical and structural information to calculate mutational effects. However, the structural input is a monomeric structure without any cofactor, ligand, ion or nucleic acids. Therefore, one should be careful for interpretation of ESCOTT results when external factors like these play an important role. Furthermore, we should note that we limited our comparison with ProteinGym v.1.0.0 and the computational methods presented therein. Some recent methods like AlphaMissense were excluded due to three reasons (40): 1) Lack of the publicly available code 2) The aim of AlphaMissense was to calculate pathogenicity of human mutations rather than other organisms. 3) Requirement of specialized hardware such as high-end graphical processing units. Moreover, we are not aware of any deep mutational scanning dataset of MPXV proteins as of writing of this paper and therefore, we do not have a complete mutational dataset for MPXV proteins for a direct comparison with our data. Despite these limitations, we present these mutational landscapes calculated with two state-of-the-art methods due to urgency of the situation with the expectation that it will be useful to scientific and clinical community.

As demonstrated in (Fig. 1b and c), ESCOTT algorithm can predict accurately impact of both single and multiple point mutations for viral proteins. Due to sheer number of double, triple or quadruple mutant combinations, we presented only single point mutational landscapes of 171 MPXV proteins in this work. Even though we have only 3 experiments containing multiple point mutations, prediction performance of ESCOTT for multiple point mutations is quite impressive. Interested researchers can conduct multiple point mutation analysis for MPXV proteins using our docker image or webserver.

Although we present a comprehensive analysis for only MPXV proteins, the approach we present here is general and computationally efficient. It can be applied very quickly to any emerging viral threat and provide valuable information on the proteins. Mutational landscapes can give information about impact of emerging variants of the virus. AMMS analysis can help to determine sensitive locations in viral proteins that can be exploited for drug design. AGMS analysis can accelerate target selection procedure against viral infections.

## Declarations

### Ethics approval and consent to participate

Not applicable. This study does not contain any individual person’s data, identifiable images, or personal information that would require any consent.

### Availability of data and material

#### Data Availability

All data generated for this study is available at https://doi.org/10.5281/zenodo.13365666.

For each one of the MPXV proteins, we have the following files:

1. Raw ESCOTT scores data file
2. Raw ESCOTT scores image in png format
3. Ranksorted ESCOTT scores data file
4. Ranksorted ESCOTT scores image in png format
5. PDB file containing pLDDT values in Bfactor column and AMMS values in occupancy column.

Each folder for a protein contains iGEMME results, which are ESCOTT calculations without structural and physico-chemical information.

Mutational landscape of each protein and their AMMS scores can be visualized using the Google Colab at https://colab.research.google.com/github/tozaktas/mpxv-mutations/blob/master/monkeypox_mutations.ipynb.

#### Code and Program Availability

ESCOTT source code is publicly available at http://gitlab.lcqb.upmc.fr/tekpinar/PRESCOTT. ESCOTT can be executed with the docker image at https://hub.docker.com/r/tekpinar/prescott-docker. Using a docker image is more appropriate for batch runs. If only mutational analyses of a few proteins are needed, we recommend ESCOTT webserver implementation at http://prescott.lcqb.upmc.fr/.

### Authors’ contributions

T.O. and A.D.T. collected the data. T.O., A.D.T. and M.T. analyzed the data. A.C. and M.T. wrote the manuscript and revised the text. All authors reviewed the manuscript.

## Acknowledgements

The numerical calculations reported in this paper were partially performed at TUBITAK ULAKBIM, High Performance and Grid Computing Center (TRUBA resources). We would like to thank them for providing us these excellent computational resources and service.

## References

1. Karagoz, A., Tombuloglu, H., Alsaeed, M., Tombuloglu, G., AlRubaish, A. A., Mahmoud, A., Smajlovic, S., Cordic, S., Rabaan, A. A. and Alsuhaimi, E. (2023) title. J Infect Public Health 16, 531–541.

2. Giorgi, F. M., Pozzobon, D., Di Meglio, A. and Mercatelli, D. (2024) title. Vaccine 42, 1841–1849.

3. Yu, Z., Zou, X., Deng, Z., Zhao, M., Gu, C., Fu, L., Xiao, W., He, M., He, L., Yang, Q., Liang, S., Wen, C. and Lu, M. (2024) title. Genomics 116, 110763.

4. Zhang, S., Li, Y. D., Cai, Y. R., Kang, X. P., Feng, Y., Li, Y. C., Chen, Y. H., Li, J., Bao, L. L. and Jiang, T. (2024) title. Frontiers in genetics 15, 1361952.

5. Chakraborty, C., Bhattacharya, M., Sharma, A. R. and Dhama, K. (2022) title. Geroscience 44, 2895–2911.

6. Tekpinar, M., David, L., Henry, T. and Carbone, A. (2025) title. Genome Biology 26, 113.

7. Mariani, V., Biasini, M., Barbato, A. and Schwede, T. (2013) title. Bioinformatics 29, 2722–2728.

8. Guo, H. B., Perminov, A., Bekele, S., Kedziora, G., Farajollahi, S., Varaljay, V., Hinkle, K., Molinero, V., Meister, K., Hung, C., Dennis, P., Kelley-Loughnane, N. and Berry, R. (2022) title. Sci Rep 12, 10696.

9. Mirdita, M., Schütze, K., Moriwaki, Y., Heo, L., Ovchinnikov, S. and Steinegger, M. (2022) title. Nat. Meth. 19, 679–682.

10. Jumper, J., Evans, R., Pritzel, A., Green, T., Figurnov, M., Ronneberger, O., Tunyasuvunakool, K., Bates, R., Zidek, A., Potapenko, A., Bridgland, A., Meyer, C., Kohl, S. A. A., Ballard, A. J., Cowie, A., Romera-Paredes, B., Nikolov, S., Jain, R., Adler, J., Back, T., Petersen, S., Reiman, D., Clancy, E., Zielinski, M., Steinegger, M., Pacholska, M., Berghammer, T., Bodenstein, S., Silver, D., Vinyals, O., Senior, A. W., Kavukcuoglu, K., Kohli, P. and Hassabis, D. (2021) title. Nature.

11. Steinegger, M., Meier, M., Mirdita, M., Vohringer, H., Haunsberger, S. J. and Soding, J. (2019) title. BMC Bioinformatics 20, 473.

12. Schrodinger, LLC (2010) The PyMOL Molecular Graphics System, Version 1.3r1.

13. Kabsch, W. and Sander, C. (1983) title. Biopolymers 22, 2577–2637.

14. Kunzmann, P. and Hamacher, K. (2018) title. BMC Bioinformatics 19, 346.

15. Saladi, S. M., Maggiolo, A. O., Radford, K. and Clemons, W. M. (2020) title. bioRxiv, 2020.2009.2022.308593.

16. Laine, E., Karami, Y. and Carbone, A. (2019) title. Mol. Biol. Evol.

17. Parigger, L., Krassnigg, A., Grabuschnig, S., Gruber, K., Steinkellner, G. and Gruber, C. C. (2023) title. Microbiol Spectr 11, e0231523.

18. Notin, P., Van Niekerk, L., Kollasch, A. W., Ritter, D., Gal, Y. and Marks, D. S. (2022) title. bioRxiv, 2022.2012.2007.519495.

19. Minasov, G., Inniss, N. L., Shuvalova, L., Anderson, W. F. and Satchell, K. J. F. (2022) title. Acta Crystallogr F Struct Biol Commun 78, 371–377.

20. Wang, X., Ma, L., Li, N. and Gao, N. (2023) title. Mol. Cell 83, 4398-4412.e4394.

21. Berman, H. M., Westbrook, J., Feng, Z., Gilliland, G., Bhat, T. N., Weissig, H., Shindyalov, I. N. and Bourne, P. E. (2000) title. Nucleic Acids Res 28, 235–242.

22. Dougherty, W. G. and Semler, B. L. (1993) title. Microbiol Rev 57, 781–822.

23. Marintcheva, B. and Weller, S. K. (2001) title. Prog. Nucleic Acid Res. Mol. Biol. 70, 77–118.

24. Guo, Y. and Yan, R. (2025) title. FEBS J. 292, 510–518.

25. Zhang, W. Z., Liu, Y. S., Yang, M. Q., Yang, J., Shao, Z. W., Gao, Y. Q., Jiang, X. R., Cui, R. X., Zhang, Y. X., Zhao, X., Shao, Q. Y., Cao, C. L., Li, H. L., Li, L. X., Liu, H. H., Gao, H. S. and Gan, J. H. (2024) title. Cell Discov 10.

26. Satheshkumar, P. S. and Moss, B. (2009) title. J. Virol. 83, 12822–12832.

27. Laliberte, J. P., Weisberg, A. S. and Moss, B. (2011) title. PLoS Pathog 7, e1002446.

28. He, J., Huang, H., Li, B., Li, H., Zhao, Y., Li, Y., Ye, W., Qi, W., Tang, W. and Wang, L. (2022) title. Front Microbiol 13, 862205.

29. Chen, N. H., Li, G. Y., Liszewski, M. K., Atkinson, J. P., Jahrling, P. B., Feng, Z. H., Schriewer, J., Buck, C., Wang, C. L., Lefkowitz, E. J., Esposito, J. J., Harms, T., Damon, I. K., Roper, R. L., Upton, C. and Buller, R. M. L. (2005) title. Virology 340, 46–63.

30. Ferrareze, P. A. G., Pereira, E. C. R. A. and Thompson, C. E. (2023) title. Arch. Virol. 168, 278.

31. Cassetti, M. C., Merchlinsky, M., Wolffe, E. J., Weisberg, A. S. and Moss, B. (1998) title. J Virol 72, 5769–5780.

32. Albarnaz, J. D., Ren, H., Torres, A. A., Shmeleva, E. V., Melo, C. A., Bannister, A. J., Brember, M. P., Chung, B. Y. and Smith, G. L. (2022) title. Nat Microbiol 7, 154–168.

33. Teissier, E., Zandomeneghi, G., Loquet, A., Lavillette, D., Lavergne, J. P., Montserret, R., Cosset, F. L., Bockmann, A., Meier, B. H., Penin, F. and Pecheur, E. I. (2011) title. PloS one 6, e15874.

34. Fischer, W. B. (2007) Viral Membrane Proteins: Structure, Function, and Drug Design, Springer Science & Business Media.

35. Chiu, W. L. and Chang, W. (2002) title. J. Virol. 76, 9575–9587.

36. Szajner, P., Jaffe, H., Weisberg, A. S. and Moss, B. (2004) title. Virology 330, 447–459.

37. Wolffe, E. J., Weisberg, A. S. and Moss, B. (1998) title. Virology 244, 20–26.

38. Boyd, O., Turner, P. C., Moyer, R. W., Condit, R. C. and Moussatche, N. (2010) title. Virology 399, 201–211.

39. Condit, R. C. and Moussatche, N. (2015) title. Virology 482, 147–156.

40. Cheng, J., Novati, G., Pan, J., Bycroft, C., Zemgulyte, A., Applebaum, T., Pritzel, A., Wong, L. H., Zielinski, M., Sargeant, T., Schneider, R. G., Senior, A. W., Jumper, J., Hassabis, D., Kohli, P. and Avsec, Z. (2023) title. Science 381, eadg7492.

