## Supplementary Information for "Comprehensive Mutational Landscape Analysis of Monkeypox Virus Proteome"

**This Supplementary Information contains 5 figures and 2 tables.**

FIGURES

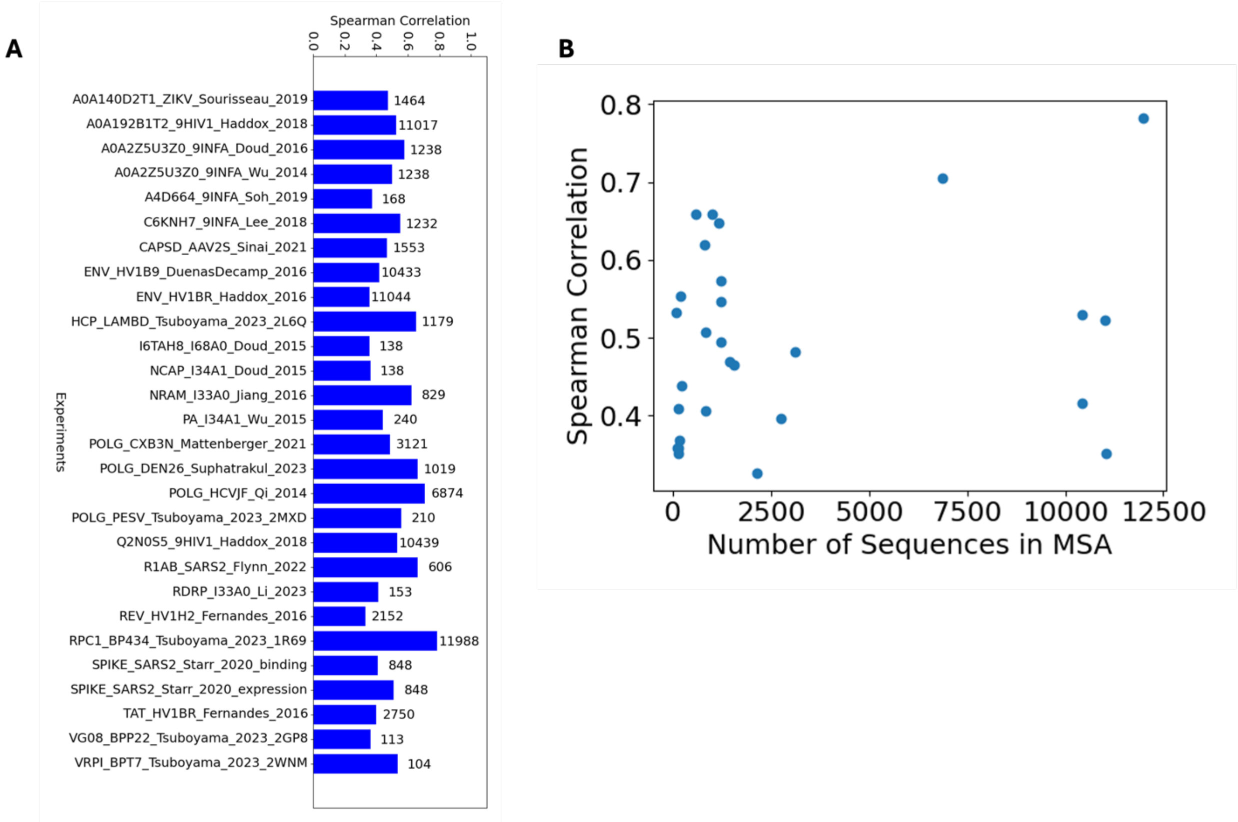


**Fig. S1** Spearman correlation and number of sequences in MSA file for 28 viral protein experiments in ProteinGym (v1.0.0. A) Bar plot of Spearman correlation for each experiment. Numbers on the right-hand side of each bar indicates number of sequences in the MSA file. B) Scatter plot of number of sequences vs Spearman correlation.

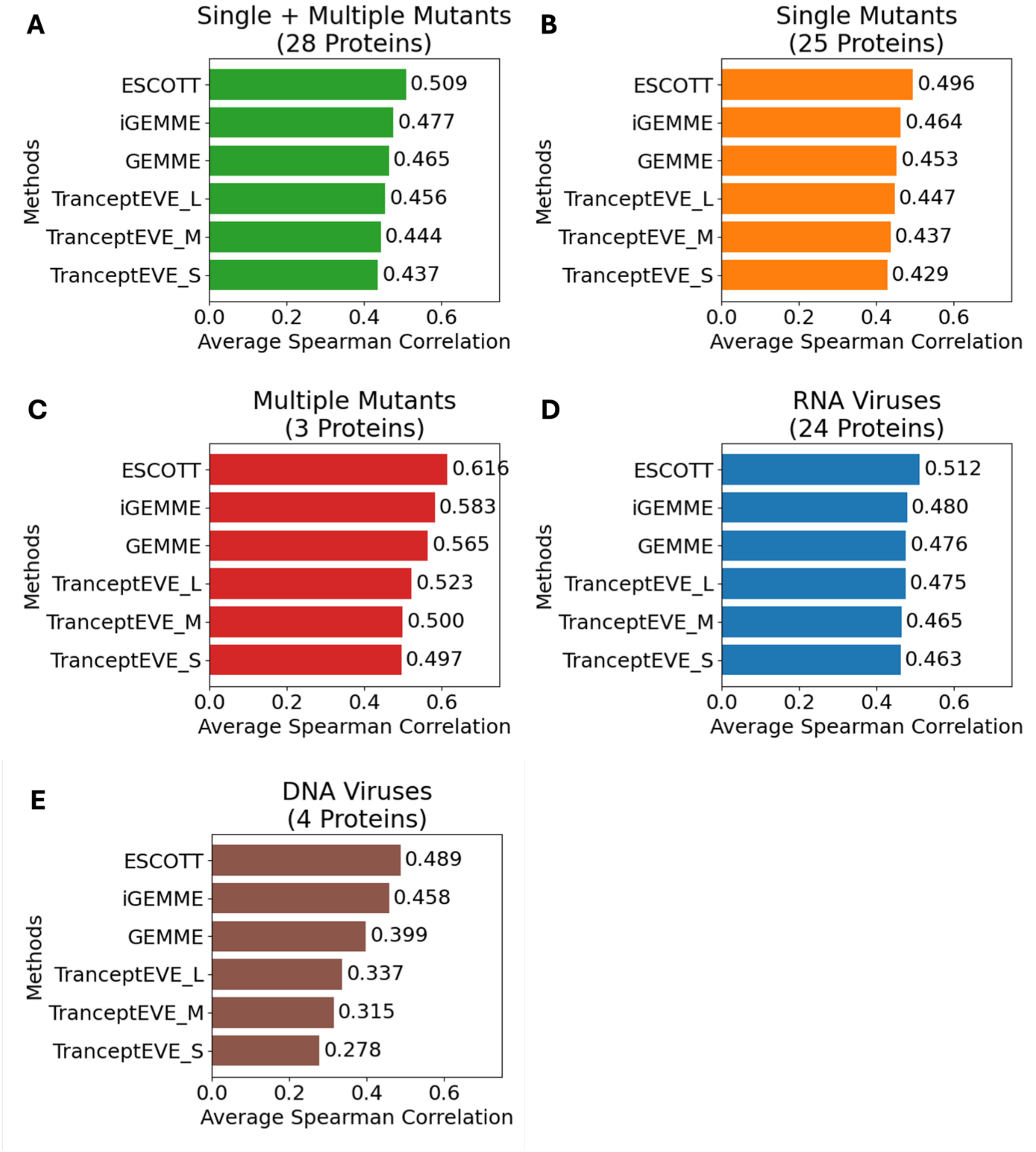


**Fig. S2** Performance comparison of ESCOTT and iGEMME with the other four methods. MSA files from ProteinGym v1.0.0 repository were used for ESCOTT and iGEMME calculations. A) Average Spearman correlation performance of the best six methods over 28 deep mutational scanning experiments of viral proteins from ProteinGym v1.0.0 dataset containing both single and multiple point mutations. B) Average Spearman correlation performance of the best six methods over 25 deep mutational scanning experiments of viral proteins from the same dataset containing only single point mutations. C) Average Spearman correlation performance of the best six methods over 3 deep mutational scanning experiments of viral proteins containing only multiple point mutations. D) Average Spearman correlation performance for proteins obtained from 24 RNA viruses. E) Average Spearman correlation performance for proteins obtained from 4 DNA viruses.

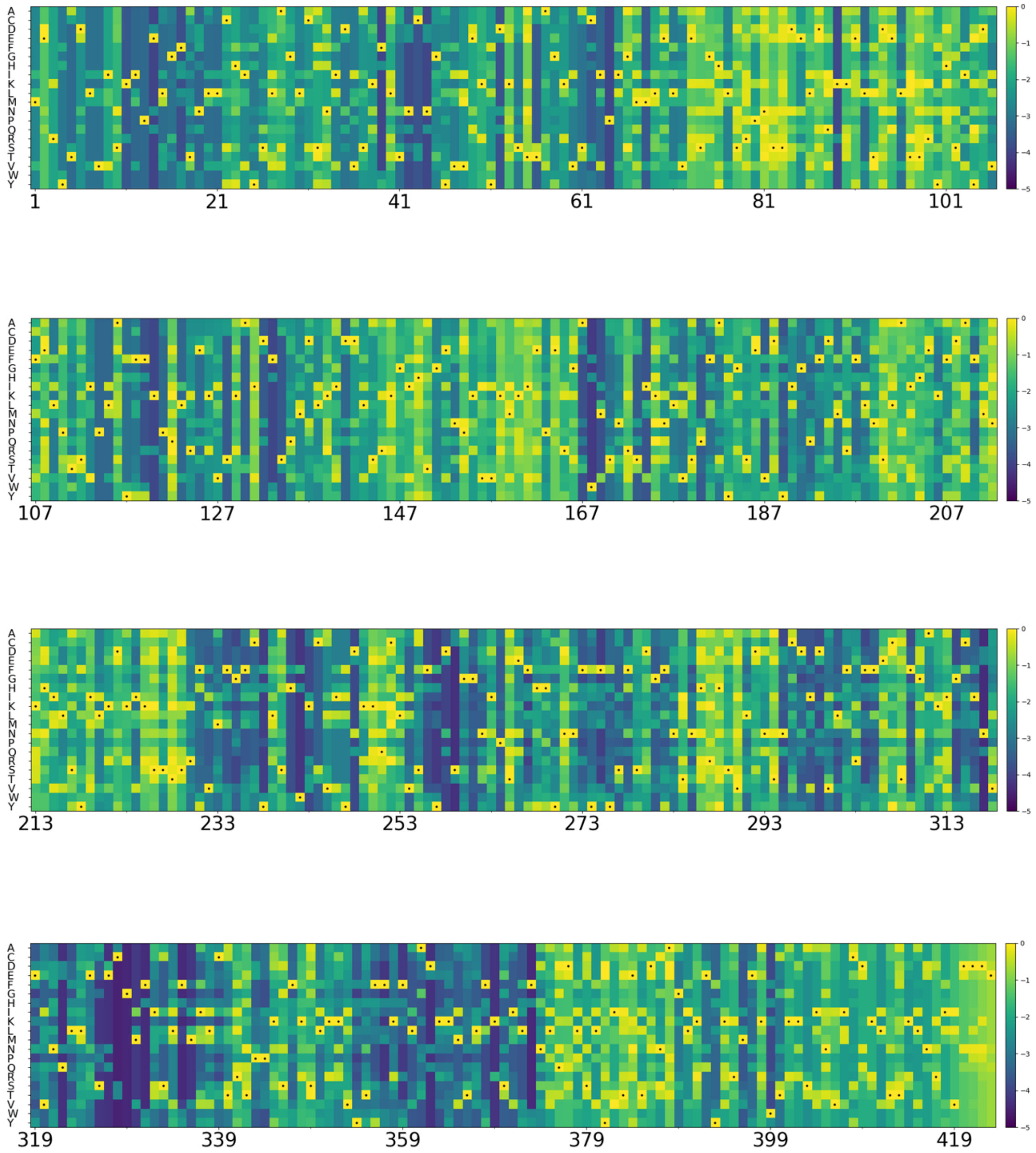


**Fig. S3** Single point mutational landscape of virus core cysteine proteinase (OPG083) calculated with ESCOTT given in raw scale and viridis colormap. Yellow color (values close to zero) indicates no effect while dark blue (values close to minus five) indicates a high effect for the mutations. Dotted squares denote the original amino acid in the protein sequence.

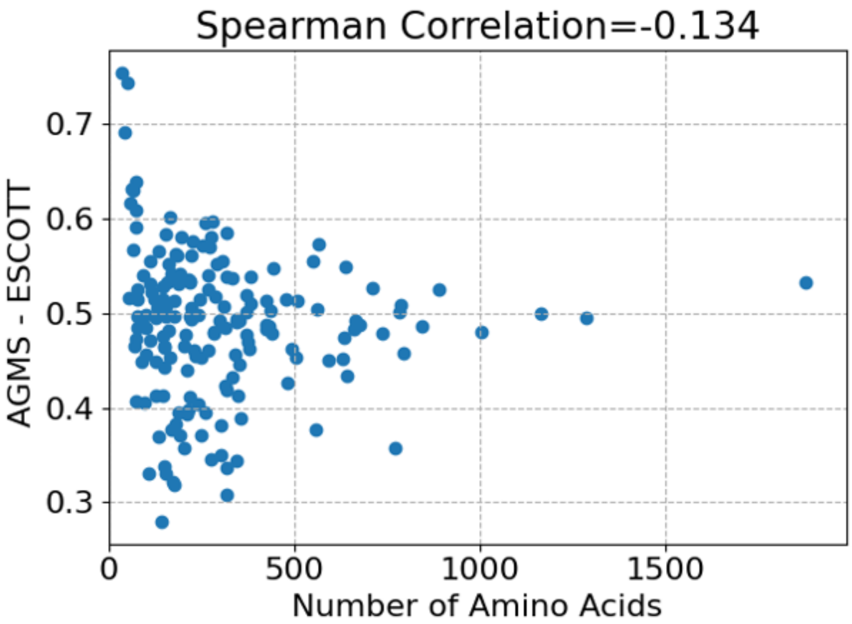


**Fig. S4** Relation between gene length and AGMS for ESCOTT. Spearman correlation between the two variables is -0.134.

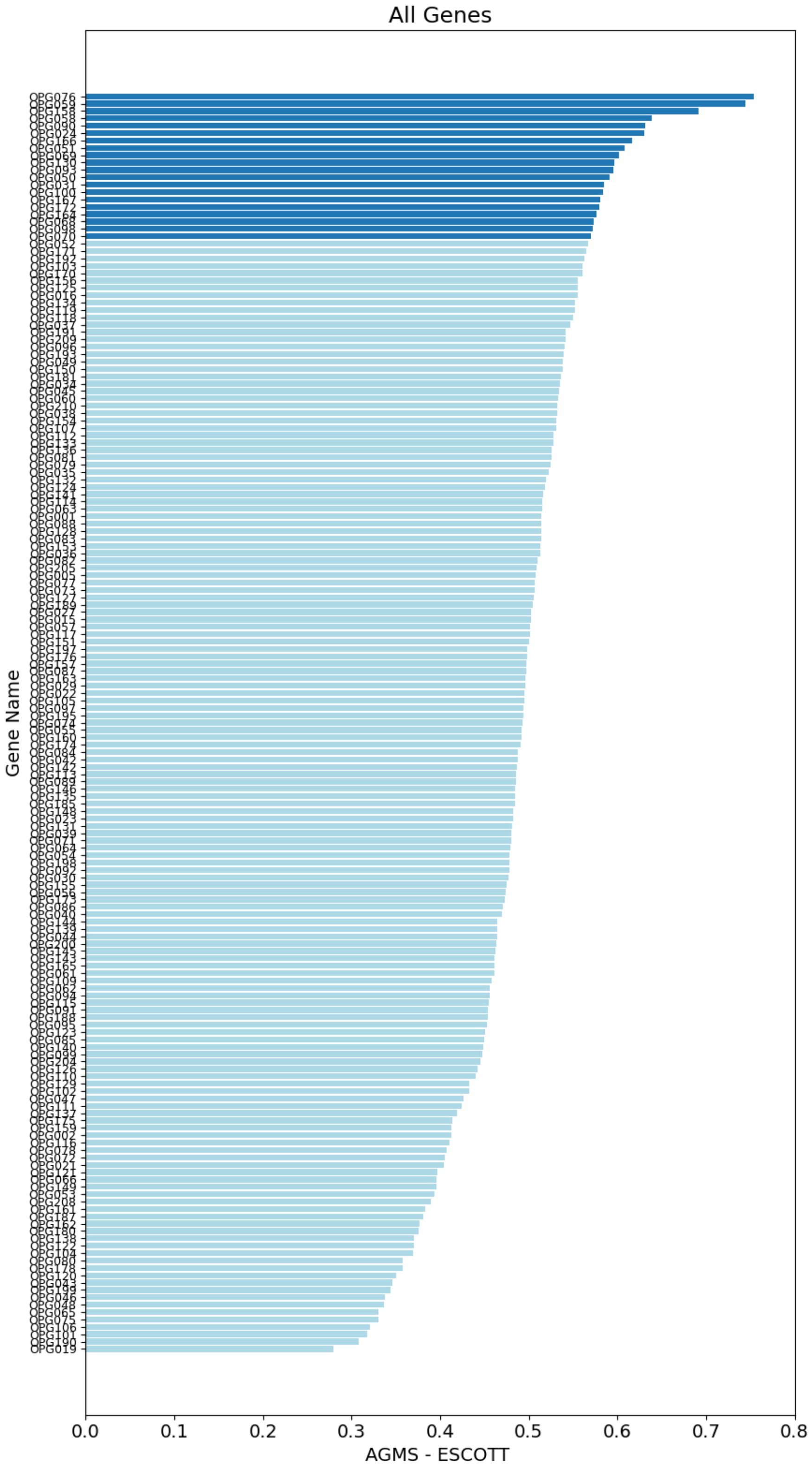


**Fig. S5** AGMS calculated from ESCOTT results for 171 proteins in MPXV genome. The top 20 proteins are colored in tab-blue and the others are colored in light-blue.

### TABLES

Table S1. The list of 171 monkeypox genes. Gene products, protein IDs, lengths of each protein and number of sequences in each multiple sequence alignment file are provided for each gene.

| Gene | Product | Protein ID | Length | # of sequences |
|---|---|---|---|---|
| **OPG001** | Chemokine binding protein | XCT41315.1 | 246 | 53 |
| **OPG002** | Crm-B secreted TNF-alpha-receptor-like protein | XCT41316.1 | 349 | 6048 |
| **OPG005** | Bcl-2-like protein | XCT41488.1 | 153 | 65 |
| **OPG015** | Ankyrin repeat protein (39) | XCT41318.1 | 437 | 2032 |
| **OPG016** | Brix domain protein | XCT41489.1 | 113 | 26 |
| **OPG019** | EGF-like domain protein | XCT41319.1 | 142 | 4885 |
| **OPG021** | Zinc finger-like protein (2) | XCT41320.1 | 242 | 8876 |
| **OPG022** | Interleukin-18-binding protein | XCT41321.1 | 126 | 246 |
| **OPG023** | Ankyrin repeat protein (2) | XCT41322.1 | 660 | 3043 |
| **OPG024** | retroviral pseudoprotease-like protein | XCT41323.1 | 64 | 16 |
| **OPG027** | Host range protein | XCT41325.1 | 150 | 80 |
| **OPG029** | Bcl-2-like protein | XCT41326.1 | 155 | 70 |
| **OPG030** | Kelch-like protein (1) | XCT41327.1 | 206 | 489 |
| **OPG031** | C4L/C10L-like family protein | XCT41328.1 | 316 | 104 |
| **OPG034** | Bcl-2-like protein | XCT41329.1 | 214 | 24 |
| **OPG035** | Bcl-2-like protein | XCT41330.1 | 117 | 53 |
| **OPG036** | Bcl-2-like protein | XCT41331.1 | 177 | 79 |
| **OPG037** | Ankyrin-like protein (1) | XCT41332.1 | 442 | 275 |
| **OPG038** | NFkB inhibitor | XCT41333.1 | 220 | 49 |
| **OPG039** | Ankyrin-like protein (3) | XCT41334.1 | 284 | 1592 |
| **OPG040** | Serpin | XCT41335.1 | 375 | 6396 |
| **OPG042** | Phospholipase-D-like protein | XCT41336.1 | 424 | 3907 |
| **OPG043** | Putative monoglyceride lipase | XCT41337.1 | 276 | 8191 |
| **OPG044** | Bcl-2-like protein | XCT41338.1 | 149 | 103 |
| **OPG045** | Caspase-9 inhibitor | XCT41339.1 | 219 | 34 |
| **OPG046** | dUTPase | XCT41340.1 | 151 | 13503 |
| **OPG047** | Kelch-like protein (2) | XCT41341.1 | 482 | 1785 |
| **OPG048** | Ribonucleotide reductase small subunit | XCT41342.1 | 319 | 11067 |
| **OPG049** | Telomere-binding protein I6 (1) | XCT41343.1 | 318 | 30 |
| **OPG050** | CPXV053 protein | XCT41344.1 | 75 | 26 |
| **OPG051** | CPXV054 protein | XCT41345.1 | 74 | 44 |
| **OPG052** | Cytoplasmic protein | XCT41346.1 | 64 | 19 |
| **OPG053** | IMV membrane protein L1R | XCT41347.1 | 212 | 140 |
| **OPG054** | Serine/threonine-protein kinase | XCT41348.1 | 439 | 175 |
| **OPG055** | Protein F11 | XCT41349.1 | 354 | 78 |
| **OPG056** | EEV maturation protein | XCT41350.1 | 635 | 84 |
| **OPG057** | Palmytilated EEV membrane protein | XCT41351.1 | 372 | 785 |
| **OPG058** | Protein F14 (1) | XCT41352.1 | 73 | 33 |
| **OPG059** | Cytochrome C oxidase | XCT41353.1 | 49 | 16 |
| **OPG060** | Protein F15 | XCT41354.1 | 158 | 48 |
| **OPG061** | Protein F16 (1) | XCT41355.1 | 231 | 136 |
| **OPG062** | DNA-binding phosphoprotein (1) | XCT41356.1 | 101 | 63 |

| | | | | |
|---|---|---|---|---|
| **OPG063** | Poly(A) polymerase catalytic subunit (3) | XCT41357.1 | 479 | 74 |
| **OPG064** | Iev morphogenesis protein | XCT41358.1 | 737 | 123 |
| **OPG065** | Double-stranded RNA binding protein | XCT41359.1 | 153 | 5869 |
| **OPG066** | DNA-directed RNA polymerase 30 kDa polypeptide | XCT41360.1 | 259 | 4798 |
| **OPG068** | IMV membrane protein E6 | XCT41361.1 | 567 | 57 |
| **OPG069** | Myristoylated protein E7 | XCT41362.1 | 166 | 43 |
| **OPG070** | Membrane protein E8 | XCT41363.1 | 273 | 52 |
| **OPG071** | DNA polymerase (2) | XCT41364.1 | 1006 | 1421 |
| **OPG072** | Sulfhydryl oxidase | XCT41365.1 | 95 | 349 |
| **OPG073** | Virion core protein E11 | XCT41366.1 | 129 | 53 |
| **OPG074** | Iev morphogenesis protein | XCT41367.1 | 665 | 90 |
| **OPG075** | Glutaredoxin-1 | XCT41368.1 | 108 | 6936 |
| **OPG076** | MV membrane EFC component | XCT41369.1 | 35 | 18 |
| **OPG077** | Telomere-binding protein I1 | XCT41370.1 | 312 | 50 |
| **OPG078** | IMV membrane protein I2 | XCT41371.1 | 73 | 41 |
| **OPG079** | DNA-binding phosphoprotein (2) | XCT41372.1 | 269 | 76 |
| **OPG080** | Ribonucleoside-diphosphate reductase (2) | XCT41373.1 | 771 | 18805 |
| **OPG081** | IMV membrane protein I5 | XCT41374.1 | 79 | 56 |
| **OPG082** | Telomere-binding protein | XCT41375.1 | 382 | 64 |
| **OPG083** | Viral core cysteine proteinase | XCT41376.1 | 423 | 116 |
| **OPG084** | RNA helicase NPH-II (2) | XCT41377.1 | 676 | 1324 |
| **OPG085** | Metalloendopeptidase | XCT41378.1 | 591 | 794 |
| **OPG086** | Entry/fusion complex component | XCT41379.1 | 111 | 59 |
| **OPG087** | Late transcription elongation factor | XCT41380.1 | 220 | 63 |
| **OPG088** | Glutaredoxin-2 | XCT41381.1 | 124 | 49 |
| **OPG089** | FEN1-like nuclease | XCT41382.1 | 434 | 94 |
| **OPG090** | DNA-directed RNA polymerase 7 kDa subunit | XCT41383.1 | 63 | 45 |
| **OPG091** | Nlpc/p60 superfamily protein | XCT41384.1 | 165 | 1410 |
| **OPG092** | Assembly protein G7 | XCT41385.1 | 371 | 67 |
| **OPG093** | Late transcription factor VLTF-1 | XCT41386.1 | 260 | 40 |
| **OPG094** | Myristylated protein | XCT41387.1 | 340 | 103 |
| **OPG095** | IMV membrane protein L1R | XCT41388.1 | 250 | 104 |
| **OPG096** | Crescent membrane and immature virion formation protein | XCT41389.1 | 92 | 46 |
| **OPG097** | Internal virion L3/FP4 protein | XCT41390.1 | 344 | 85 |
| **OPG098** | Nucleic acid binding protein VP8/L4R | XCT41391.1 | 251 | 51 |
| **OPG099** | Membrane protein CL5 | XCT41392.1 | 128 | 67 |
| **OPG100** | IMV membrane protein J1 | XCT41393.1 | 152 | 55 |
| **OPG101** | Thymidine kinase | XCT41394.1 | 177 | 8819 |
| **OPG102** | Cap-specific mRNA | XCT41395.1 | 333 | 208 |
| **OPG103** | DNA-directed RNA polymerase subunit | XCT41396.1 | 185 | 54 |
| **OPG104** | Myristylated protein | XCT41397.1 | 133 | 188 |
| **OPG105** | DNA-dependent RNA polymerase subunit rpo147 | XCT41398.1 | 1286 | 1907 |
| **OPG106** | Tyr/ser protein phosphatase | XCT41399.1 | 171 | 4923 |
| **OPG107** | Entry-fusion complex essential component | XCT41400.1 | 189 | 82 |
| **OPG109** | RNA polymerase-associated transcription-specificity factor RAP94 | XCT41402.1 | 795 | 91 |
| **OPG110** | Late transcription factor VLTF-4 (1) | XCT41403.1 | 210 | 83 |
| **OPG111** | DNA topoisomerase type I | XCT41404.1 | 314 | 745 |

| | | | | |
|---|---|---|---|---|
| **OPG112** | Late protein H7 | XCT41405.1 | 146 | 61 |
| **OPG113** | mRNA capping enzyme large subunit | XCT41406.1 | 845 | 190 |
| **OPG114** | Virion protein D2 | XCT41407.1 | 146 | 75 |
| **OPG115** | Virion core protein D3 | XCT41408.1 | 233 | 67 |
| **OPG116** | Uracil DNA glycosylase superfamily | XCT41409.1 | 218 | 929 |
| **OPG117** | NTPase (1) | XCT41410.1 | 785 | 129 |
| **OPG118** | Early transcription factor 70 kDa subunit | XCT41411.1 | 637 | 556 |
| **OPG119** | RNA polymerase subunit RPO18 | XCT41412.1 | 161 | 50 |
| **OPG120** | Carbonic anhydrase | XCT41413.1 | 304 | 6842 |
| **OPG121** | NUDIX domain protein | XCT41414.1 | 213 | 5118 |
| **OPG122** | MutT motif protein | XCT41415.1 | 248 | 6433 |
| **OPG123** | Nucleoside triphosphatase I | XCT41416.1 | 631 | 1310 |
| **OPG124** | mRNA capping enzyme small subunit | XCT41417.1 | 287 | 66 |
| **OPG125** | Rifampicin resistance protein | XCT41418.1 | 551 | 84 |
| **OPG126** | Late transcription factor VLTF-2 (2) | XCT41419.1 | 150 | 75 |
| **OPG127** | Late transcription factor VLTF-3 (1) | XCT41420.1 | 224 | 80 |
| **OPG128** | S-S bond formation pathway protein | XCT41421.1 | 77 | 60 |
| **OPG129** | Virion core protein P4b | XCT41422.1 | 644 | 101 |
| **OPG130** | A5L protein-like | XCT41423.1 | 281 | 35 |
| **OPG131** | DNA-directed RNA polymerase 19 kDa subunit | XCT41424.1 | 161 | 73 |
| **OPG132** | Virion morphogenesis protein | XCT41425.1 | 372 | 62 |
| **OPG133** | Early transcription factor 82 kDa subunit | XCT41426.1 | 710 | 195 |
| **OPG134** | Intermediate transcription factor VITF-3 (1) | XCT41427.1 | 292 | 61 |
| **OPG135** | IMV membrane protein A9 | XCT41428.1 | 100 | 76 |
| **OPG136** | Virion core protein P4a | XCT41429.1 | 891 | 88 |
| **OPG137** | Viral membrane formation protein | XCT41430.1 | 318 | 60 |
| **OPG138** | A12 protein | XCT41431.1 | 190 | 93 |
| **OPG139** | IMV membrane protein A13L | XCT41432.1 | 70 | 16 |
| **OPG140** | IMV membrane protein A14 | XCT41433.1 | 90 | 56 |
| **OPG141** | DUF1029 domain protein | XCT41434.1 | 53 | 42 |
| **OPG142** | Core protein A15 | XCT41435.1 | 94 | 44 |
| **OPG143** | Myristylated protein | XCT41436.1 | 377 | 99 |
| **OPG144** | IMV membrane protein P21 | XCT41437.1 | 204 | 69 |
| **OPG145** | DNA helicase | XCT41438.1 | 492 | 273 |
| **OPG146** | Zinc finger-like protein (1) | XCT41439.1 | 77 | 67 |
| **OPG148** | DNA polymerase processivity factor | XCT41441.1 | 426 | 60 |
| **OPG149** | Holliday junction resolvase | XCT41442.1 | 187 | 369 |
| **OPG150** | Intermediate transcription factor VITF-3 (2) | XCT41443.1 | 382 | 71 |
| **OPG151** | DNA-dependent RNA polymerase subunit rpo132 | XCT41444.1 | 1164 | 3666 |
| **OPG153** | Orthopoxvirus A26L/A30L protein | XCT41445.1 | 509 | 91 |
| **OPG154** | IMV surface fusion protein | XCT41446.1 | 110 | 103 |
| **OPG155** | Envelope protein A28 homolog | XCT41447.1 | 146 | 72 |
| **OPG156** | DNA-directed RNA polymerase 35 kDa subunit | XCT41448.1 | 305 | 67 |
| **OPG157** | IMV membrane protein A30 | XCT41449.1 | 78 | 54 |
| **OPG158** | A32.5L | XCT41450.1 | 42 | 15 |
| **OPG159** | CPXV166 protein | XCT41451.1 | 145 | 46 |
| **OPG160** | ATPase A32 | XCT41452.1 | 300 | 265 |

| | | | | |
|---|---|---|---|---|
| **OPG161** | EEV glycoprotein (1) | XCT41453.1 | 181 | 1084 |
| **OPG162** | EEV glycoprotein (2) | XCT41454.1 | 168 | 753 |
| **OPG163** | MHC class II antigen presentation inhibitor | XCT41455.1 | 176 | 60 |
| **OPG164** | IEV transmembrane phosphoprotein | XCT41456.1 | 228 | 36 |
| **OPG165** | CPXV173 protein | XCT41457.1 | 268 | 68 |
| **OPG166** | hypothetical protein | XCT41458.1 | 59 | 30 |
| **OPG167** | CD47-like protein | XCT41459.1 | 277 | 245 |
| **OPG170** | Chemokine binding protein | XCT41460.1 | 221 | 74 |
| **OPG171** | Profilin domain protein | XCT41461.1 | 133 | 215 |
| **OPG172** | Type-I membrane glycoprotein | XCT41462.1 | 196 | 23 |
| **OPG173** | MPXVgp154 | XCT41463.1 | 74 | 24 |
| **OPG174** | Hydroxysteroid dehydrogenase | XCT41464.1 | 346 | 2516 |
| **OPG175** | Copper/zinc superoxide dismutase | XCT41465.1 | 125 | 6216 |
| **OPG176** | Bcl-2-like protein | XCT41466.1 | 240 | 45 |
| **OPG178** | Thymidylate kinase | XCT41467.1 | 204 | 7702 |
| **OPG180** | DNA ligase (2) | XCT41468.1 | 559 | 6709 |
| **OPG181** | M137R | XCT41469.1 | 334 | 51 |
| **OPG185** | Hemagglutinin | XCT41470.1 | 313 | 777 |
| **OPG187** | Ser/thr kinase | XCT41471.1 | 303 | 3710 |
| **OPG188** | Schlafen (1) | XCT41472.1 | 503 | 1440 |
| **OPG189** | Ankyrin repeat protein (25) | XCT41473.1 | 561 | 1204 |
| **OPG190** | EEV type-I membrane glycoprotein | XCT41474.1 | 317 | 8979 |
| **OPG191** | Ankyrin-like protein (46) | XCT41475.1 | 168 | 36 |
| **OPG192** | Virulence protein | XCT41476.1 | 182 | 16 |
| **OPG193** | Soluble interferon-gamma receptor-like protein | XCT41477.1 | 267 | 68 |
| **OPG195** | Intracellular viral protein | XCT41478.1 | 221 | 38 |
| **OPG197** | CPXV205 protein | XCT41479.1 | 100 | 27 |
| **OPG198** | Ser/thr kinase | XCT41480.1 | 282 | 1877 |
| **OPG199** | Serpin | XCT41481.1 | 344 | 9633 |
| **OPG200** | Bcl-2-like protein | XCT41482.1 | 149 | 165 |
| **OPG204** | IFN-alpha/beta-receptor-like secreted glycoprotein | XCT41483.1 | 351 | 429 |
| **OPG205** | Ankyrin repeat protein (44) | XCT41484.1 | 787 | 2002 |
| **OPG208** | Serpin | XCT41485.1 | 357 | 9293 |
| **OPG209** | Virulence protein | XCT41486.1 | 190 | 39 |
| **OPG210** | B22R family protein | XCT41487.1 | 1880 | 177 |

Table S2. Deep mutational scanning experiments of viral proteins from ProteinGym v1.0.0 and their Spearman correlations with results of 6 prediction algorithms. Experiments highlighted with red text color contain multiple point mutations. Four DNA virus proteins are highlighted with bold font.

| DMS_id | ESCOTT | iGEMME | GEMME | TranceptEVE_L | TranceptEVE_M | TranceptEVE_S | Number of Mutants | Genetic Material | Number of Sequences |
|---|---|---|---|---|---|---|---|---|---|
| A0A140D2T1_ZIKV_Sourisseau_2019 | 0.469 | 0.442 | 0.430 | 0.373 | 0.358 | 0.361 | 9576 | RNA | 1464 |
| A0A192B1T2_9HIV1_Haddox_2018 | 0.523 | 0.510 | 0.496 | 0.528 | 0.518 | 0.524 | 12577 | RNA | 11017 |
| A0A2Z5U3Z0_9INFA_Doud_2016 | 0.573 | 0.552 | 0.538 | 0.573 | 0.576 | 0.557 | 10715 | RNA | 1238 |
| A0A2Z5U3Z0_9INFA_Wu_2014 | 0.495 | 0.485 | 0.513 | 0.558 | 0.556 | 0.541 | 2350 | RNA | 1238 |
| A4D664_9INFA_Soh_2019 | 0.369 | 0.337 | 0.443 | 0.461 | 0.439 | 0.434 | 14421 | RNA | 168 |
| C6KNH7_9INFA_Lee_2018 | 0.547 | 0.518 | 0.475 | 0.452 | 0.441 | 0.439 | 10754 | RNA | 1232 |
| **CAPSD_AAV2S_Sinai_2021** | **0.465** | **0.542** | **0.445** | **0.426** | **0.343** | **0.338** | **42328** | **DNA** | **1553** |
| ENV_HV1B9_DuenasDecamp_2016 | 0.416 | 0.400 | 0.389 | 0.392 | 0.392 | 0.390 | 375 | RNA | 10433 |
| ENV_HV1BR_Haddox_2016 | 0.352 | 0.342 | 0.350 | 0.365 | 0.369 | 0.367 | 12863 | RNA | 11044 |
| **HCP_LAMBD_Tsuboyama_2023_2L6Q** | **0.647** | **0.635** | **0.579** | **0.507** | **0.430** | **0.374** | **1040** | **DNA** | **1179** |
| I6TAH8_I68A0_Doud_2015 | 0.352 | 0.296 | 0.368 | 0.401 | 0.399 | 0.383 | 9462 | RNA | 138 |
| NCAP_I34A1_Doud_2015 | 0.359 | 0.291 | 0.377 | 0.441 | 0.426 | 0.425 | 9462 | RNA | 138 |
| NRAM_I33A0_Jiang_2016 | 0.619 | 0.576 | 0.635 | 0.632 | 0.638 | 0.628 | 298 | RNA | 829 |
| PA_I34A1_Wu_2015 | 0.439 | 0.331 | 0.584 | 0.584 | 0.592 | 0.588 | 1820 | RNA | 240 |
| POLG_CXB3N_Mattenberger_2021 | 0.482 | 0.481 | 0.495 | 0.458 | 0.430 | 0.387 | 15711 | RNA | 3121 |
| POLG_DEN26_Suphatrakul_2023 | 0.659 | 0.624 | 0.622 | 0.563 | 0.397 | 0.469 | 16897 | RNA | 1019 |
| POLG_HCVJF_Qi_2014 | 0.705 | 0.673 | 0.630 | 0.560 | 0.528 | 0.480 | 1630 | RNA | 6874 |
| POLG_PESV_Tsuboyama_2023_2MXD | 0.554 | 0.414 | 0.507 | 0.452 | 0.463 | 0.453 | 5130 | RNA | 210 |
| Q2N0S5_9HIV1_Haddox_2018 | 0.530 | 0.517 | 0.507 | 0.513 | 0.515 | 0.526 | 12729 | RNA | 10439 |
| R1AB_SARS2_Flynn_2022 | 0.659 | 0.632 | 0.558 | 0.565 | 0.567 | 0.546 | 5725 | RNA | 606 |
| RDRP_I33A0_Li_2023 | 0.409 | 0.391 | 0.520 | 0.526 | 0.508 | 0.494 | 12003 | RNA | 153 |
| REV_HV1H2_Fernandes_2016 | 0.327 | 0.315 | 0.282 | 0.235 | 0.261 | 0.246 | 2147 | RNA | 2152 |
| RPC1_BP434_Tsuboyama_2023_1R69 | 0.782 | 0.768 | 0.743 | 0.690 | 0.693 | 0.699 | 1459 | RNA | 11988 |
| SPIKE_SARS2_Starr_2020_binding | 0.406 | 0.287 | 0.247 | 0.341 | 0.346 | 0.356 | 3802 | RNA | 848 |
| SPIKE_SARS2_Starr_2020_expression | 0.507 | 0.451 | 0.359 | 0.475 | 0.455 | 0.451 | 3798 | RNA | 848 |
| TAT_HV1BR_Fernandes_2016 | 0.396 | 0.380 | 0.368 | 0.268 | 0.295 | 0.366 | 1577 | RNA | 2750 |
| **VG08_BPP22_Tsuboyama_2023_2GP8** | **0.359** | **0.270** | **0.487** | **0.409** | **0.403** | **0.336** | **723** | **DNA** | **113** |
| **VRPI_BPT7_Tsuboyama_2023_2WNM** | **0.532** | **0.438** | **0.084** | **0.006** | **0.083** | **0.064** | **1047** | **DNA** | **104** |
